# Dual-Channel Event Microscopy for Ultrafast Biological Imaging

**DOI:** 10.1101/2025.11.01.685972

**Authors:** Ruipeng Guo, Xueli Pan, Qilin Deng, Abrar Ahmed, Qianwan Yang, Joseph Greene, Tongyu Li, Suet Ying Chan, Zhixiong Chen, Guorong Hu, Hui Feng, Lei Tian

## Abstract

Many fundamental biological processes—spanning immune–tumor interactions, neuronal signaling, and microvascular flow—exhibit fast, multiscale dynamics among diverse cell types within three-dimensional tissue environments. Capturing such activity requires imaging systems that simultaneously achieve high temporal resolution, multicolor capability, and volumetric coverage over large fields of view (FOVs). However, existing modalities remain limited by trade-offs among imaging speed, spectral capacity, depth of field (DOF), and spatial resolution. Here, we present Dual-Channel Event Microscopy (DEM), which integrates digital micromirror device (DMD)–based pulsed illumination, extended-DOF optics, and event-based sensing for ultrafast, dual-channel volumetric imaging across a 2.3 × 1.3 mm^2^ FOV with an effective 200 *µ*m DOF. Using dual-color fluorescent phantoms and microsphere flow assays, DEM achieves accurate spectral separation and reconstruction of rapid motion at kilohertz frame rates. *In vivo*, DEM enables simultaneous visualization of neutrophils and premalignant tumors in freely swimming zebrafish. In immobilized specimens, it provides robust, sensor-level optical sectioning near the heart, suppressing diffuse background to reveal fine vascular networks and active blood circulation into and out of the cardiac chambers. DEM further enables quantitative mapping of blood-flow dynamics in the zebrafish tail, resolving arterial–venous differences and capturing heartbeat-driven oscillations that reflect cardiac pumping with high temporal fidelity. By uniting ultrafast acquisition, dual-channel capability, volumetric coverage, and intrinsic optical sectioning within a single event-driven architecture, DEM offers a powerful platform for visualizing rapid multicellular interactions and physiological dynamics in living systems.

## Introduction

Advances in live imaging have enabled researchers to probe complex biological dynamics spanning immune–tumor interactions [1, 2], neuronal signaling [3, 4], and microvascular flow [5, 6]. These processes entail fast, multiscale dynamics among diverse cell types within three-dimensional (3D) tissue environments. Capturing such phenomena requires imaging systems that combine high temporal resolution, multicolor capability, volumetric coverage, and a wide field of view (FOV). However, existing approaches remain constrained by intrinsic trade-offs among imaging speed, spectral capacity, depth of field (DOF), and spatial resolution [7, 8]. Depth coverage typically demands axial scanning [9], while multi-channel architectures often sacrifice speed and photon efficiency [10]. Conversely, systems optimized for high temporal resolution usually restrict FOV or depth coverage, or compromise spatial resolution [11].

Recent efforts have sought to overcome these constraints through ultrafast volumetric imaging [12]. Multi-z confocal microscopy [13, 14] and fast-scanning two-photon microscopy systems [15–20] can achieve kilohertz-rate volumetric recording, yet at the cost of limited FOV, increased optical complexity, and potential phototoxicity. Light-field microscopy, by capturing angular information in a single exposure within a widefield setup, enables volumetric reconstruction of sparse dynamic events, recently reaching kilohertz rates by leveraging spatially compressed column readout in squeezed light-field microscopy [21] or event-based sensing [22]. Nevertheless, the intrinsic spatial–angular multiplexing in light-field imaging imposes fundamental trade-offs between lateral resolution, depth coverage, and FOV, while reducing light throughput [23, 24]. Furthermore, these methods typically support only single-color acquisition, whereas multichannel imaging techniques—such as spectral splitting and multi-illumination approaches—operate at lower frame rates [25, 26], precluding faithful capture of ultrafast biological events. Overcoming these competing constraints within a single, versatile imaging platform remains a central challenge.

Here, we present Dual-channel Event Microscopy (DEM), a versatile platform for ultrafast, dual-channel volumetric fluorescence imaging across large FOVs. At its core, DEM employs an event camera in which each pixel operates independently and asynchronously, reporting local intensity changes with microsecond temporal precision. This architecture eliminates global exposure cycles and enables immediate response to transient signals, allowing biological dynamics to be captured at their native timescales while minimizing redundant data acquisition [27]. The result is a substantial improvement in temporal responsiveness and data efficiency compared with conventional frame-based or scanning systems [22].

To achieve dual-channel imaging, DEM integrates DMD-modulated illumination with asynchronous event sensing. The DMD operates in a kilohertz-rate binary mode, where alternating on/off micromirror states sequentially excite two spectral channels, encoding color information in time without sacrificing speed or photon efficiency (Fig. 1A). The resulting spectrally distinct fluorescence signals are detected by a single event camera (Fig. 1B). Each DMD switching cycle induces rapid intensity transitions that generate events on the sensor, enabling faithful capture of both dynamic and static signals. The event camera’s native latency—on the order of several hundred microseconds—is comparable to the DMD flip rate, allowing temporal encoding of spectral information without color filters. Consequently, dual-channel imaging speed is governed directly by the DMD switching frequency.

**Figure 1:**
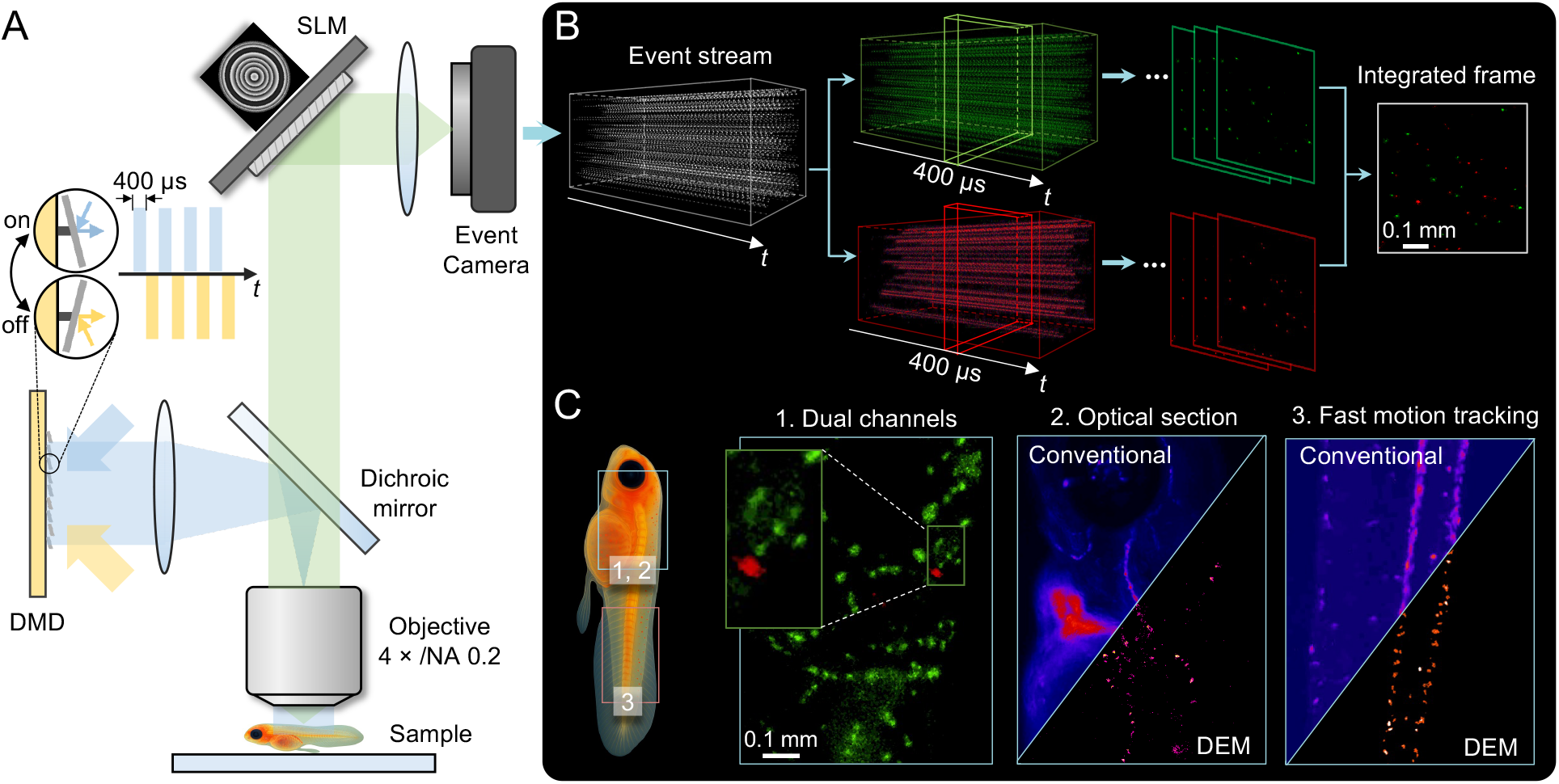
Dual-Channel Event Microscopy (DEM) for ultrafast, extended-DOF biological imaging. (A) Setup: A DMD alternates excitation between two spectral channels, while an SLM encodes EDOF in the pupil plane. Fluorescence is combined by a dichroic mirror and recorded by an event camera for kilohertz-rate, dual-channel imaging. (B) Processing: The event stream is temporally separated by DMD switching, reconstructed into channel-specific frames, and integrated to form composite dual-color images. (C) In vivo imaging: (1) Dual-channel visualization of neutrophils (green) and premalignant tumor cells (red) in zebrafish. (2) Comparison of conventional and event-based imaging near the heart shows enhanced optical sectioning and vascular contrast with DEM. (3) Tail-region comparison highlights DEM’s ability to track GFP-labeled red blood cells without motion artifacts.

To enable single-shot volumetric imaging, DEM uses SLM-based pupil engineering to extend the DOF. A custom phase mask displayed on the SLM generates an elongated point spread function (PSF) that maintains focus across hundreds of micrometers, resulting in more than a tenfold DOF extension. The same phase mask is jointly optimized for both spectral channels to align their focal planes and ensure a consistent DOF response in dual-color imaging. This approach optically compresses volumetric information into a single 2D projection while preserving spatial fidelity across the extended DOF (EDOF) range.

DEM enables dual-channel imaging at kilohertz rates and achieves an EDOF exceeding 200 *µ*m for both channels across a 2.3 × 1.3 mm^2^ FOV, supporting volumetric imaging over large spatial scales. Notably, in samples with strong background scattering, DEM inherently achieves optical sectioning by detecting only dynamic, in-focus intensity changes (Fig. 1C). Because events are triggered solely by temporal contrast, temporally slowly varying diffuse and static background photons remain unregistered, providing sensor-level sectioning without the need for confocal apertures or structured illumination. This selective sensitivity enhances image contrast and SNR in complex biological environments, enabling better visualization of dynamic structures in scattering tissues.

We first validated DEM using 3D dual-color fluorescent phantoms and microsphere flow assays, demonstrating both accurate spectral separation and faithful reconstruction of rapid volumetric motion at kilohertz rates. We then applied DEM *in vivo* to freely swimming zebrafish, a particularly challenging setting due to their rapid, unrestrained motion. The microsecond responsiveness of the event sensor enabled real-time tracking of fast activity, preserving cellular detail and spatial fidelity even during whole-body movement. Dual-color imaging of labeled neutrophils and premalignant tumor cells revealed localized multicellular interactions within the moving organism, illustrating DEM’s capacity to probe complex dynamics in natural behavioral states. In immobilized zebrafish, DEM further demonstrated complementary advantages in both spatial and temporal performance. Around the cardiac region, where conventional sCMOS-based widefield fluorescence imaging is degraded by out-of-focus background and scattering, DEM achieved sensor-level optical sectioning by detecting only temporally varying signals under static illumination. This selective response effectively suppressed diffuse fluorescence and revealed clear blood circulation into and out of the heart, resolving fine vascular pathways and dynamic flow patterns with enhanced contrast and clarity (Fig. 1C). In the tail region without strong background and scattering, DEM enabled quantitative mapping of blood-flow dynamics, resolving arterial–venous differences and capturing heartbeat-driven oscillations under pulsed illumination (Fig. 1C). Together, these results highlight DEM’s unique ability to deliver motion-robust, dual-color, and temporally precise imaging across spatial scales—extending fluorescence microscopy into regimes where motion, scattering, and complexity have traditionally imposed severe limits.

We establish DEM as a high-speed, dual-channel fluorescence imaging platform that overcomes long-standing limitations of ultrafast microscopy—restricted FOV, depth coverage, and single-color operation. By integrating DMD-based illumination, SLM-enabled extended DOF optics, and asynchronous event sensing within a single architecture, DEM jointly optimizes temporal, spectral, and volumetric performance to achieve kilohertz-rate imaging across extended depths. This co-designed illumination–optics–sensor framework provides an efficient, information-rich representation of dynamic biological processes. More broadly, DEM represents a new paradigm in microscopy—one that leverages event-driven acquisition to adaptively match the dynamics of living systems, enabling scalable and information-efficient observation of multicellular processes in motion.

## Result

### Dual-channel Event Microscopy system

The working principle of DEM is illustrated in Fig. 1. Illumination from blue and lime LEDs is directed onto the DMD at incident angles of +24^*°*^and –24^*°*^relative to the surface normal. Operating in a kilohertz-rate binary mode, the DMD toggles its micromirrors between ±12^*°*^to alternately excite the two spectral channels. This rapid switching merges both beams into a shared optical path while providing precise temporal separation between colors. The on and off mirror states correspond to alternating blue and lime illumination (Fig. 1A). The resulting spectrally distinct fluorescence signals are routed through a dual-band optical path, modulated in the Fourier plane by an SLM carrying an engineered phase mask for EDOF imaging, and finally detected by a single event camera (Fig. 1B).

Each DMD switching cycle introduces abrupt illumination transitions that generate intensity changes readily detected by the event camera. The native event latency—on the order of several hundred microseconds—is well matched to the DMD’s micromirror switching rate (up to 22.7 kHz), allowing efficient capture of rapid excitation changes. Pixel-level characterization using a uniform fluorescent slide revealed asymmetric response dynamics between positive and negative events (SI Appendix, Fig. S2): positive events, corresponding to brightness increments, responded more rapidly, whereas negative events exhibited slower recovery due to charge decay following illumination transitions. The latency for both event types decreased sharply with increasing excitation power, reaching ∼200 *µ*s under illumination conditions comparable to those used in biological imaging.

Although the DMD and event camera operate asynchronously, the periodic illumination modulation introduces a stable temporal signature that is directly reflected in the event timestamps. Consequently, spectral information is encoded in time, enabling the entire pixel array to capture both channels without color filters. Individual pixels independently report red- or green-channel events, which are later separated by timestamp during reconstruction. Experimental validation confirms that the temporally interleaved channels exhibit minimal crosstalk and can be reconstructed into continuous dual-color video streams. This illumination-modulation strategy achieves kilohertz-rate, dual-channel imaging without compromising temporal resolution, spatial fidelity, or photon efficiency. Prior to detection, the fluorescence passes through the SLM, which displays a phase mask optimized using our previously developed genetic algorithm [28]. This mask extends the DOF by more than tenfold while preserving lateral resolution. Importantly, the same phase profile is jointly optimized for both spectral channels, aligning their focal planes and ensuring uniform depth response. The resulting EDOF configuration enables light-efficient, single-shot capture of volumetric samples, eliminating the need for mechanical axial scanning.

During post-processing, events are temporally sorted according to the DMD modulation sequence and integrated into channel-specific frames. These reconstructed frames are merged into synchronized dual-color videos, providing high-speed visualizations of dynamic biological activity.

### System characterization

We characterized the system to evaluate its resolution performance, EDOF capability, and dual-channel fidelity. Figure 2A shows PSF measurements acquired with both sCMOS and event cameras in the red and green channels. Compared with the unmodified PSF (ref), the engineered PSF exhibits an approximately tenfold axial extension, while the event camera reproduces a depth response closely matching that of the sCMOS camera. The corresponding full-width-at-half-maximum (FWHM) values within a 200 *µ*m axial range are plotted in Fig. 2B, confirming that lateral resolution is well preserved throughout the extended depth range. To further validate spatial fidelity, we imaged fluorescent resolution targets positioned at −100*µ*m, 0 *µ*m, and +100 *µ*m (Fig. 2C). Both color channels maintained high-contrast features and consistent alignment across depths, demonstrating robust EDOF performance and uniform lateral resolution.

**Figure 2:**
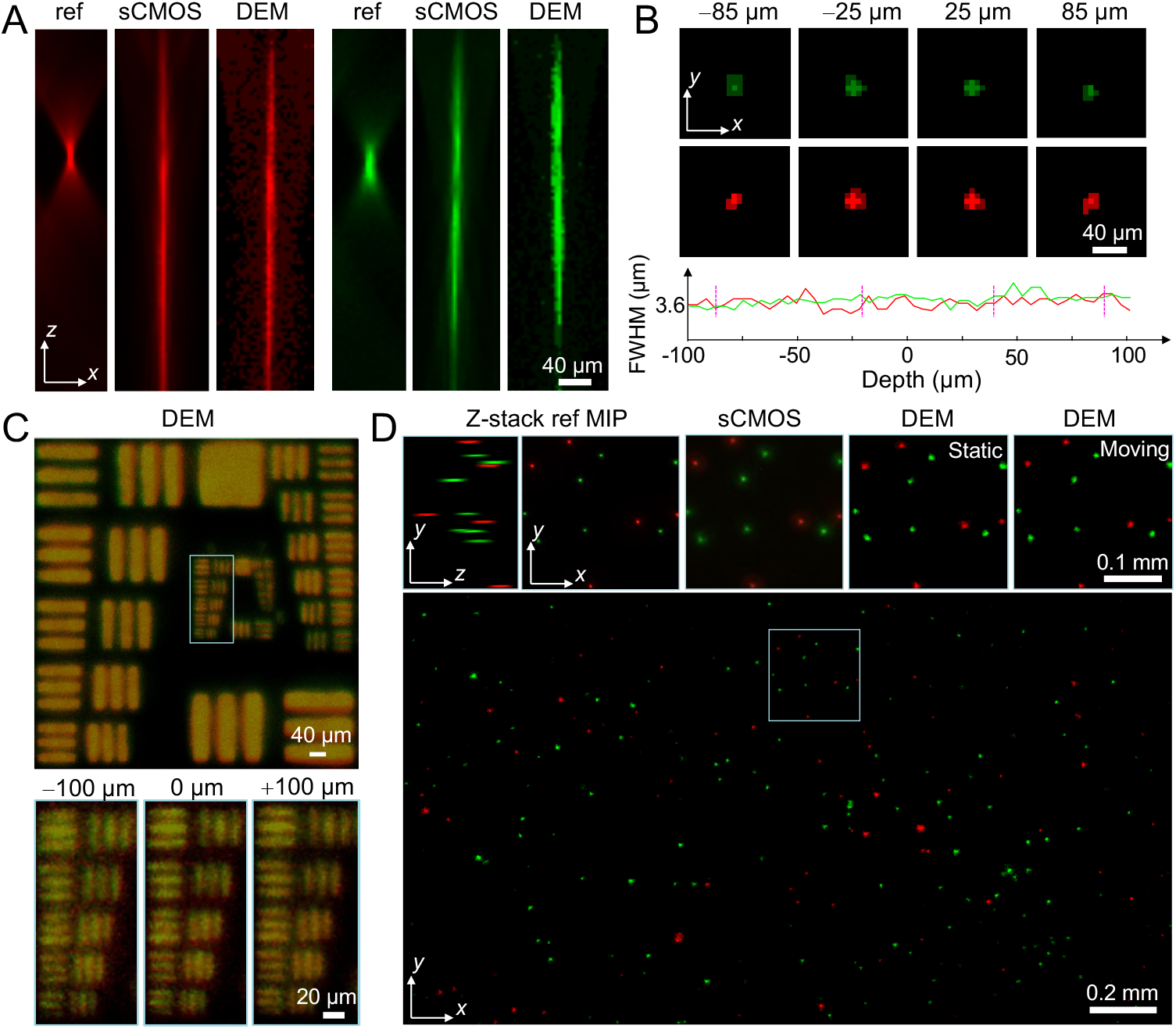
System characterization and performance validation. (A) PSF measurements for red and green fluorescence channels acquired using a reference wide-field system (ref), an sCMOS camera under EDOF encoding (sCMOS), and the event camera under the same EDOF encoding (DEM). (B) Event PSFs remain laterally consistent across the extended axial range, as confirmed by FWHM analysis over depth. (C) Imaging of fluorescent resolution targets at three axial positions (−100 *µ*m, 0 *µ*m, and +100 *µ*m) demonstrates preserved lateral resolution across depths in both channels. (D) Imaging of a dual-color fluorescent bead phantom: comparison among wide-field reference MIPs, sCMOS-based EDOF imaging, and event-based EDOF imaging for both static and moving samples, showing consistent dual-color imaging, and minimal spectral crosstalk over the extended focal volume.

Next, we evaluated DEM using a dual-color fluorescent bead phantom to assess its performance on both static and dynamic samples. The phantom, with a thickness of 200 *µ*m, was first imaged using a standard wide-field fluorescence microscope to obtain a sequentially acquired dual-color *z*-stack reference, and subsequently imaged with both the sCMOS camera (sequential excitation) and the event-based camera (simultaneous dual-channel) under identical EDOF modes. To evaluate temporal fidelity, we additionally imaged the same phantom while introducing controlled lateral motion using the event-based EDOF system. As shown in Fig. 2D, reconstructions from the event-based system closely matched the maximum-intensity projections (MIPs) of both the wide-field reference and sCMOS-based EDOF results. Notably, the event camera maintained consistent color separation with minimal spectral crosstalk, accurately distinguishing the two fluorescence channels based solely on their temporally encoded timestamps. Across the full 2.3mm × 1.3 mm FOV, the event-based system reliably captured both stationary and moving fluorescent beads, demonstrating high-speed, dual-color volumetric imaging with spectral fidelity comparable to gold-standard sequential acquisition methods.

Additional validation on a fixed brain slice is presented in SI Appendix, Fig. S5, demonstrating the performance of the dual-color event-encoded imaging under controlled, static conditions. In this test, neurons expressing eGFP and tdTomato were imaged under alternating blue and lime excitation. Independent single-channel recordings confirmed that positive and negative events correspond to sequential on/off excitation within each color channel, whereas in the dual-color configuration, the events from both channels are temporally multiplexed. The resulting composite reconstruction shows accurate channel separation and high spatial fidelity, confirming that the system achieves crosstalk-free dual-color imaging under alternating excitation.

### Kilohertz imaging of dynamic phantoms and fluorescent microsphere flow

We evaluated the system’s high-speed performance through controlled fluorescent microsphere experiments. First, a 3D phantom (200 *µ*m thick) composed of red and green microspheres was mounted on a motorized stage and translated laterally at 2.3 mm/s during acquisition. The DMD alternated excitation between spectral channels every 0.4 ms, enabling 1.25 kHz dual-color imaging. Events generated from the two channels were temporally separated and integrated into frames. Figure 3A shows the reconstructed image of the moving phantom across the entire FOV. Time-resolved snapshots (Fig. 3B) reveal spatially distinct red and green trajectories on microsecond timescales, illustrating DEM’s temporal precision. Snapshots spaced 36 ms apart align with the programmed stage motion, confirming accurate high-speed tracking.

**Figure 3:**
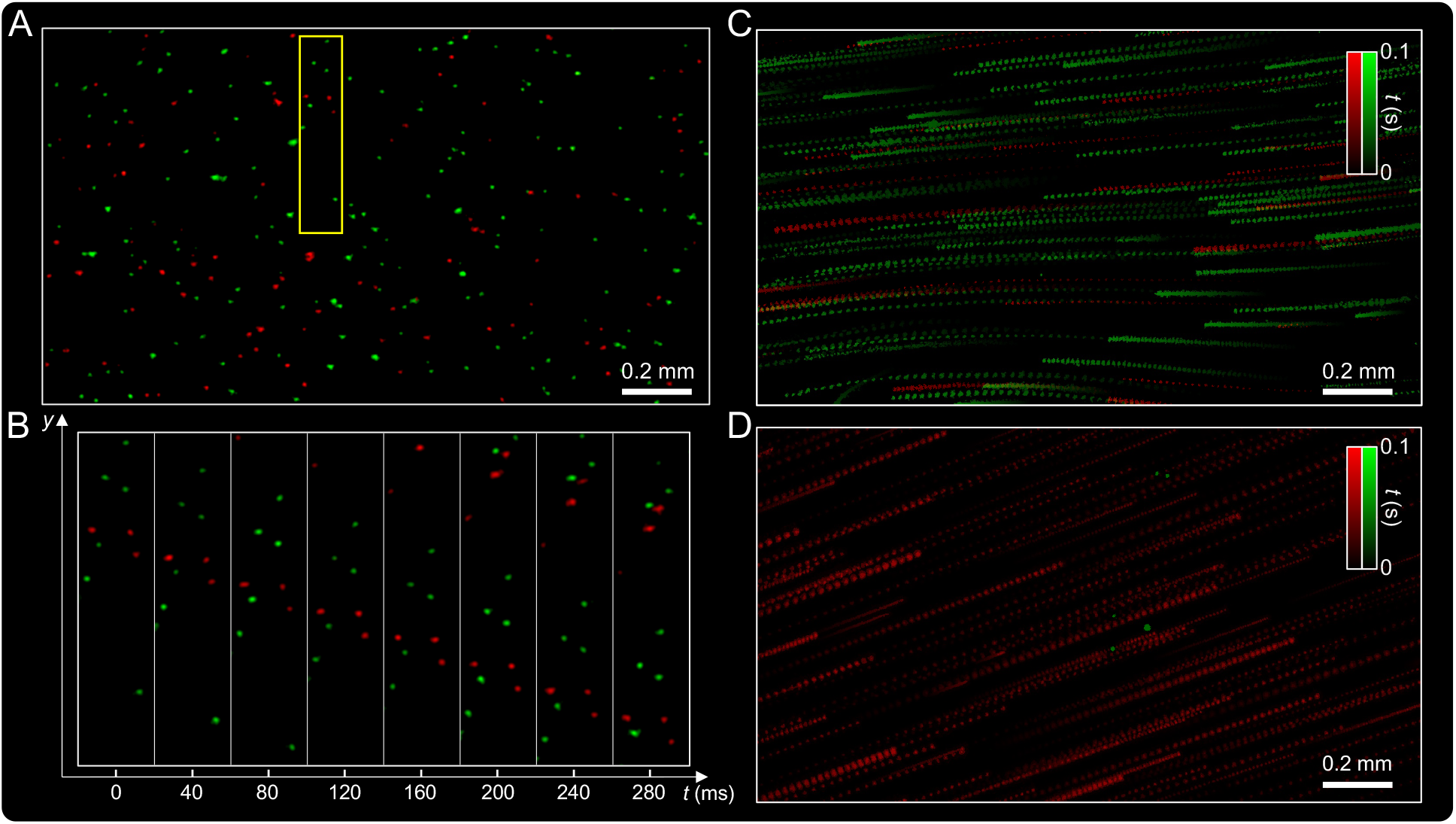
Kilohertz-rate dual-color imaging of dynamic phantom and fluorescent microsphere flow. (A) Reconstructed event-based image of a fast-moving fluorescent phantom comprising red and green microspheres, demonstrating high-speed dual-channel imaging across a 2.3 mm × 1.3 mm FOV. (B) Temporally resolved zoom-ins from the yellow box in (A), showing microsecond-scale particle motion with clear red–green separation across sequential time bins (horizontal axis: time; vertical axis: spatial displacement along the *y* axis). (C) Dual-color microsphere flow captured in a microfluidic channel, visualized with temporal color encoding to represent events over a 0.1 s acquisition. (D) Mixed-flow experiment with red microspheres in motion and green microspheres immobilized on the top and bottom channel surfaces, demonstrating DEM’s ability to distinguish dynamic and static signals in complex flow environments.

To emulate fast biological flow, we fabricated a microfluidic channel and introduced a mixed suspension of red and green fluorescent microspheres. The captured dual-color flow (Fig. 3C) is visualized using temporal color encoding to represent particle dynamics. Microspheres moving at different depths (within ∼200 *µ*m) were clearly resolved. Velocity analysis of the reconstructed frames revealed particle speeds ranging from 1 to 9 mm/s. With an integration time of 0.5 ms per channel, even the fastest particles were recorded without motion blur, corresponding to an effective dual-color frame rate of 1 kHz.

In a mixed-flow configuration, red microspheres were set in motion while green microspheres were immobilized on the top and bottom channel surfaces. As shown in Fig. 3D, DEM successfully distinguished moving and stationary particles within the same FOV, demonstrating robust dual-color imaging under complex flow conditions. Together, these experiments highlight DEM’s ability to perform high-speed tracking, resolve multiple spectral channels with minimal crosstalk, and quantify particle velocities—capabilities that enable precise analysis of microfluidic flows at kilohertz frame rates.

### Real-time tracking of freely moving Zebrafish with premalignant tumor

To assess DEM’s capability for dual-color volumetric imaging of biological dynamics *in vivo*, we imaged zebrafish bearing mCherry-labeled premalignant tumors and GFP-labeled neutrophils. The specimens measured approximately 3 mm in length and 300 *µ*m in thickness. As a reference, the head region of an anesthetized zebrafish was imaged using conventional widefield fluorescence microscopy with axial scanning and sequential dual-color illumination. Figure 4A shows MIPs in both *x*–*y* and *x*–*z* planes, revealing the 3D distribution of neutrophils and tumor cells. Using DEM, we imaged the same region and reconstructed both cell populations in a single frame (Fig. 4B). Although minor positional differences arise from cellular motions between imaging sessions, the overall correspondence of cellular features confirms reliable dual-color EDOF imaging with accurate spectral separation and spatial consistency.

**Figure 4:**
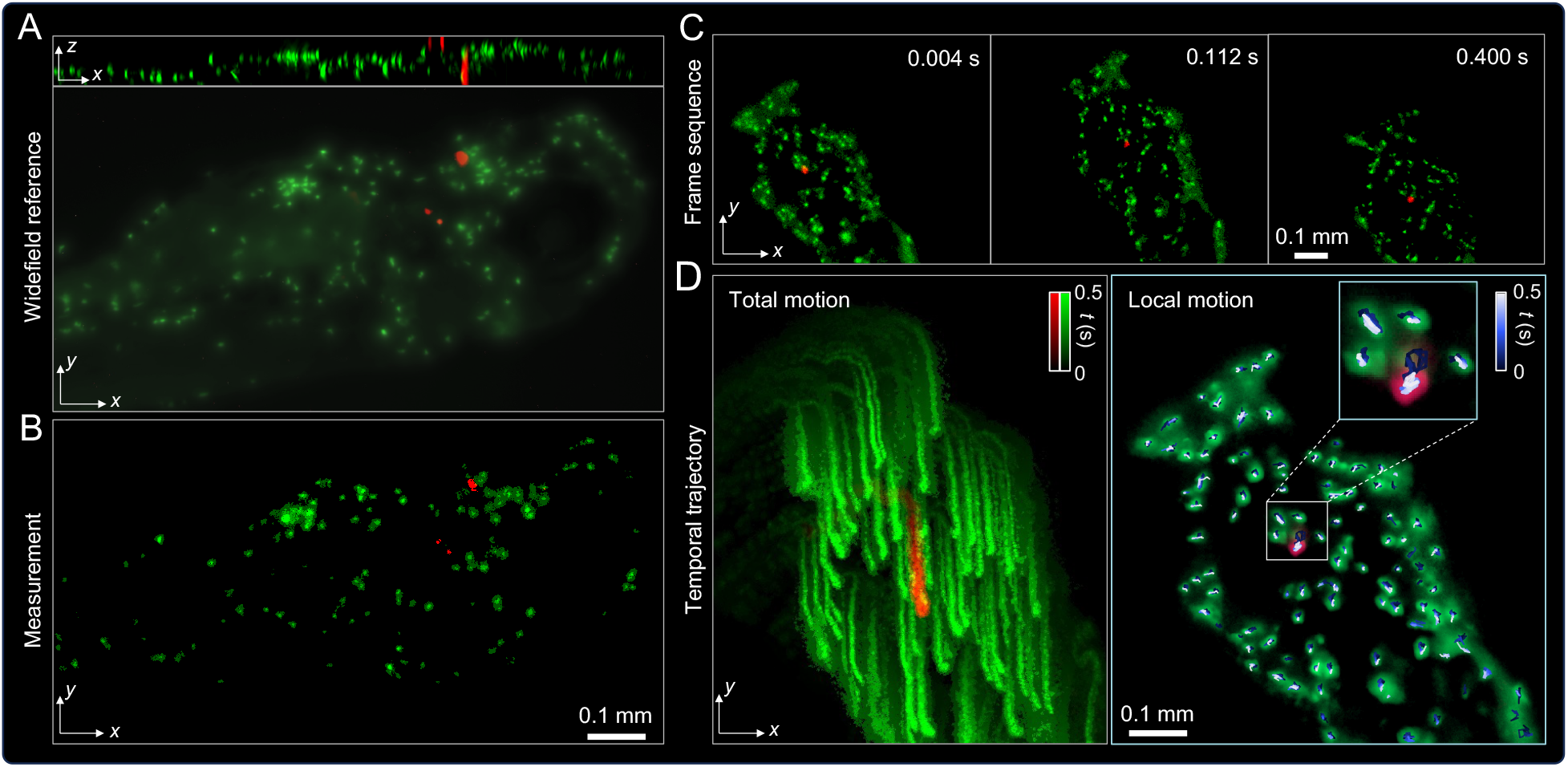
Imaging of freely swimming zebrafish with dual-color labeling of premalignant tumor and immune cells. (A) Widefield reference obtained from an axial scan showing GFP-labeled neutrophils (green) and mCherry-labeled premalignant tumor cells (red). (B) Corresponding dual-channel EDOF image acquired by DEM, simultaneously capturing tumor and immune cells in a single snapshot. (C) Time-lapse frames from high-speed DEM recording of a freely swimming zebrafish, capturing concurrent dynamics of neutrophils and tumor cells. (D) Temporally color-coded trajectories over a 0.5 s interval. Left: total motion including whole-body swimming. Right: Local cellular motion after global movement subtraction, visualized by color-coded motion tracks overlaid on the MIP of registered event frames.

We next imaged freely swimming zebrafish placed in a droplet of water on a coverslip mounted on the sample stage. As the fish moved through the FOV, DEM continuously captured high-speed, dual-channel recordings. The DMD switching interval was set to 2 ms, providing an effective dual-channel frame rate of 250 Hz—sufficient to resolve cellular dynamics during natural locomotion. Representative frames in Fig. 4C show clear visualization of both neutrophils and premalignant tumor cells at different positions during the fish’s motion. Figure 4D (left) shows a temporally color-coded MIP over a 0.5 s interval, revealing the free motion of individual cells within the zebrafish. Global body motion was removed using a rigid-body registration algorithm, isolating the local displacements of labeled cells. Compared with the unregistered stack (Total motion) in Fig. 4D (left), the averaged MIP of the registered stack (Local motion) in Fig. 4D (right) exhibits no residual global displacement, confirming effective motion stabilization. Trajectories were color-coded by acquisition time (blue → white, 0–0.5 s) to illustrate the temporal evolution of local motion within the field of view. The inset in Fig. 4D highlights representative cells and their trajectories, demonstrating local motion dynamics after global registration. Within the 0.5 s observation window, neutrophil positions remained largely stable, consistent with their intermittent motility and slow patrolling behavior at this timescale.

These results demonstrate DEM’s unique capability for high-speed, volumetric, and multiplexed fluorescence imaging in freely moving organisms. The combination of event-based sensing, EDOF, and dual-color excitation enables real-time visualization of immune–tumor interactions in vivo—opening the door to longitudinal studies of cellular behavior in unrestrained, physiologically relevant settings.

### Event-based optical sectioning and real-time visualization of blood cell dynamics in vivo

We further demonstrated DEM’s capability for optical sectioning and real-time flow imaging in live zebrafish at 4 days postfertilization, using GFP-labeled red blood cells. The specimen was positioned within the imaging FOV and illuminated via the blue LED channel. For single-color samples, illumination was configured in either constant or DMD-modulated mode, depending on the sample’s optical properties and the desired balance between temporal resolution and signal contrast.

In regions with dense fluorescence and strong background, constant illumination without EDOF was employed to assess DEM’s inherent optical sectioning capability. Under constant illumination, events are generated solely by dynamic fluorescence changes, while static and multiply scattered photons remain unregistered. Because the event stream is defined by temporal contrast rather than fixed frame exposure, the effective frame rate is determined post hoc during event integration and is practically limited by SNR. As shown in Fig. 5A, the conventional sCMOS image (left) exhibits strong background fluorescence, whereas the event-based image (right) isolates moving red blood cells with high contrast and minimal background—demonstrating sensor-level optical sectioning without confocal apertures or structured illumination.

**Figure 5:**
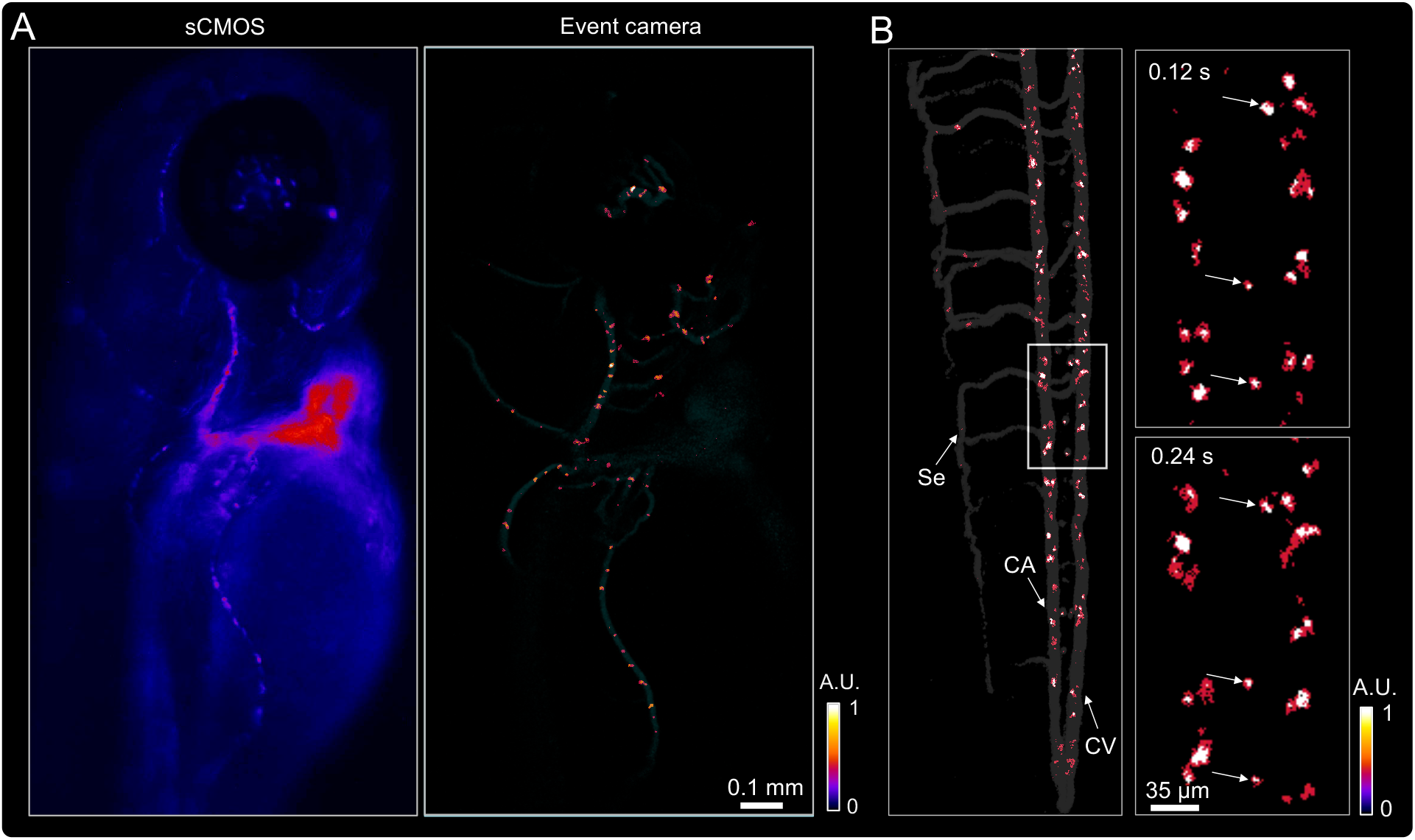
Event-based optical sectioning and real-time imaging of blood cell dynamics in live zebrafish. (A) Comparison of GFP-labeled red blood cells imaged in the zebrafish head region using a conventional sCMOS camera and the event-based system. The sCMOS image exhibits strong background fluorescence, whereas the event-based image selectively highlights in-focus, dynamic signals with minimal background. (B) Reconstructed event-based frame showing GFP-labeled blood cells in the CA, CV, and Se vessels in the tail region. Sequential zoom-ins reveal blur-free trajectories of moving cells and static cells (white arrows) across consecutive time points.

For tail-region imaging with reduced scattering, the DMD modulation was activated with a 2 ms switching interval to capture both moving and stationary cells. A single DEM reconstructed frame (Fig. 5B, left) resolves blood cells within the caudal artery (CA), caudal vein (CV), and segmental vessels (Se). Sequential zoom-ins (Fig. 5B, right) show blur-free trajectories of flowing cells alongside stationary ones labeled with white arrows, confirming high temporal fidelity. A comparison with constant-illumination imaging (SI Appendix, Fig. S4) further demonstrates that DMD modulation enables simultaneous visualization of both dynamic and static red blood cells, enhancing sensitivity while preserving temporal precision. These results demonstrate that DEM achieves both real-time optical sectioning in scattering tissue and quantitative visualization of blood flow dynamics across extended vascular networks.

### Quantification of blood flows in live zebrafish

To quantify hemodynamics, we integrated event signals over time to generate a temporal MIP, revealing the vascular architecture in the tail region (Fig.6A). Quantitative flow analysis was performed using kymographs, which plot spatial position along a vessel against time, visualizing motion as streaks whose slopes correspond to velocity. High-speed DEM reconstructions enabled extraction of kymographs along user-defined trajectories (light-blue lines in Fig.6A). As shown in Fig. 6B, in the CA and CV regions (*x*_1_–*x*_3_), diagonal streaks represent moving red blood cells, with the slope indicating flow velocity and periodic oscillations reflecting heartbeat-driven pulsation. In the Se vessel (*x*_4_), two vertical traces mark stationary cells (white arrows), while intermittent diagonal or dotted lines between them correspond to transiently moving cells. This spatiotemporal pattern underscores DEM’s ability to distinguish dynamic from static cells and to resolve directional blood flow with submillisecond temporal fidelity.

**Figure 6:**
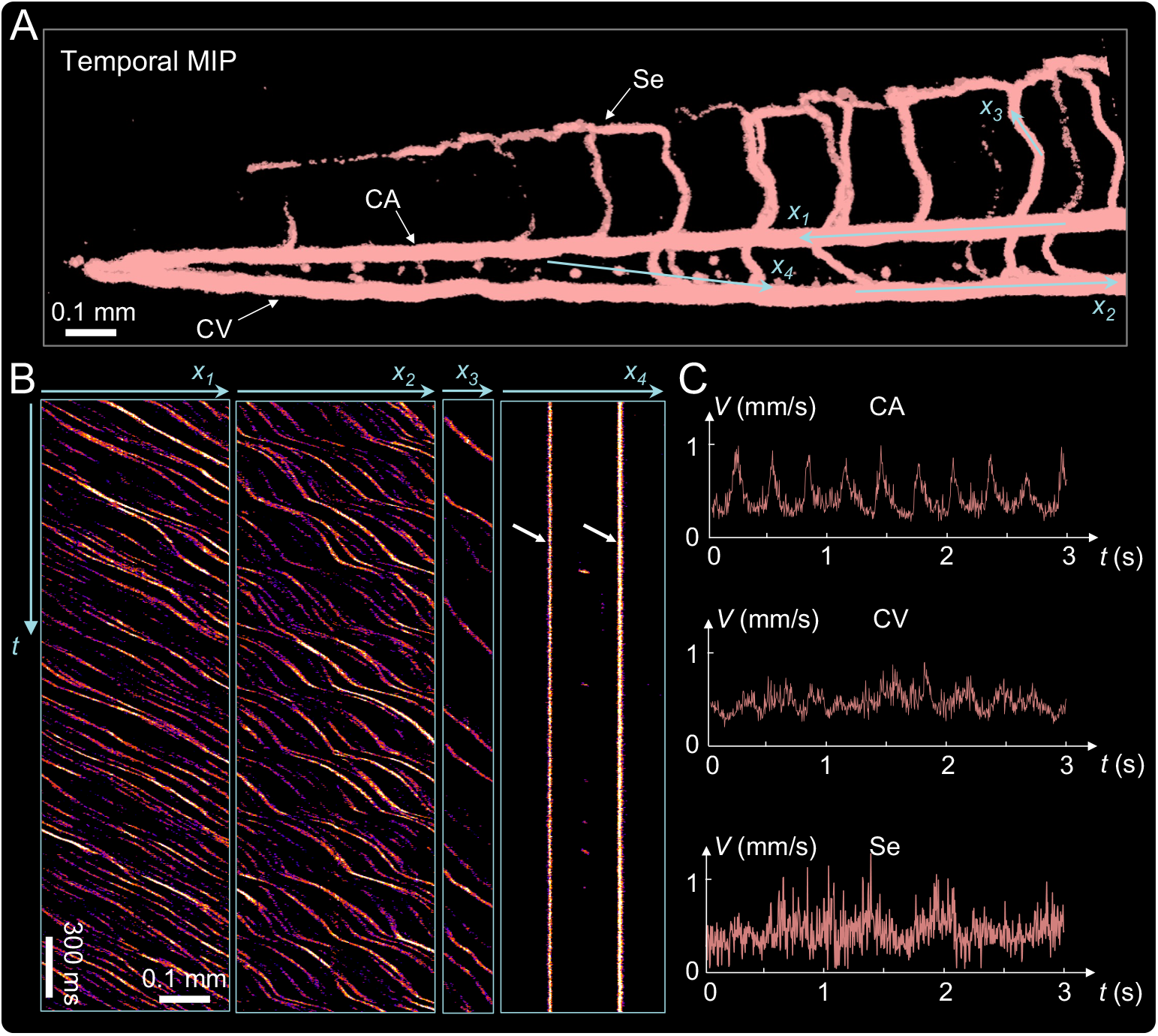
Quantification of blood cell dynamics in live zebrafish. (A) Temporal MIP of recorded events from DEM revealing the vascular network, including the CA, CV, and Se. (B) Kymographs extracted along lines in (A) illustrate time-resolved trajectories in CA, CV, and Se vessels. (C) Velocity profiles derived from temporal stacks show blood cell speeds of 0.2–0.9 mm/s, with periodic oscillations corresponding to the heartbeat and higher peak velocities in CA compared to CV and Se.

Velocity profiles derived from the temporal stacks are shown in Fig.6C. Blood cell speeds ranged from 0.2 to 0.9mm/s, with periodic fluctuations corresponding to the heartbeat. CA exhibited the highest peak velocities, while CV and Se showed slower, steadier flow, consistent with expected arterial–venous circulation patterns. These quantitative results confirm that DEM can resolve both steady and pulsatile hemodynamics with high temporal precision across multiple vessel types within the same FOV, providing a powerful tool for *in vivo* vascular analysis.

## Discussion

We developed a computational microscopy platform that integrates DMD-based time-multiplexed excitation, engineered EDOF optics, and asynchronous event detection to enable kilohertz-rate, dual-channel fluorescence imaging across large FOVs and extended depths. Through this illumi-nation–optics–sensor co-design, the system jointly optimizes temporal, spectral, and volumetric performance, allowing high-speed, motion-free visualization of complex biological dynamics without mechanical scanning or multi-camera architectures. By registering only dynamic photons, the system provides intrinsic optical sectioning in scattering tissues and accurate reconstruction of rapid processes such as blood flow and immune–tumor interactions. Together, these capabilities establish a unified framework for information-efficient, multi-channel imaging across large space–time scales, overcoming the trade-offs among speed, depth, and contrast that limit conventional frame-based microscopy.

Relative to light-field ultrafast approaches [21, 22], DEM preserves full-aperture resolution and photon efficiency while avoiding the spatial–angular trade-offs and reconstruction complexity inherent to multiplexed capture. The resulting uniformly sharp projections across extended depths are particularly advantageous for sparsely labeled biological specimens, where information-rich, motion-free visualizations provide sufficient detail for mechanistic insight and quantitative analysis [29].

Future refinements could further enhance performance through inverse- or learning-based PSF optimization [30], adaptive DMD modulation to improve photon efficiency and event triggering, and advanced denoising algorithms tailored for event-driven biological data. Such developments would expand DEM’s sensitivity and applicability to more complex biological processes.

In summary, DEM introduces a scalable framework for ultrafast, dual-channel, volumetric fluorescence imaging that adaptively matches the dynamics of living systems. Its combination of speed, optical sectioning, and multiplexing opens new opportunities for probing neural, vascular, and immune processes in their natural temporal context, advancing quantitative and mechanistic understanding across the biological sciences.

## Methods

### DEM setup

A schematic of the DEM system is shown in Fig. S1 (see SI Appendix). In the illumination module, a high-power blue LED (SOLIS-470C, Thorlabs) and a lime LED (SOLIS-565D, Thorlabs) are positioned symmetrically at incident angles of ±24^*°*^relative to the surface normal of the DMD (DLP7000, Digital Light Innovations). The emitted light from each LED is collimated by condenser lenses, filtered through band-pass excitation filters (ET475/35x, Thorlabs; ET575/35x, Chroma), and relayed onto the DMD via an additional lens to form quasi-parallel illumination at the micromirror plane. The DMD operates in a high-speed binary mode with micromirror tilt angles of ±12^*?*^, where the on and off states correspond to alternating excitation by the blue and lime LEDs. This rapid mirror switching temporally multiplexes the two illumination channels within a shared optical path, enabling synchronized dual-color excitation without mechanical or optical switching elements.

Illumination power was optimized to balance fluorescence signal intensity across channels for each experiment. For phantom and bead-flow measurements, the blue and lime channels were set to 3.2 mW mm^−2^ and 2.1 mW mm^−2^, respectively. For zebrafish with premalignant tumors, the powers were adjusted to 3.2 mW mm^−2^ (blue) and 4.2 mW mm^−2^ (lime) to equalize emission across the green and red channels. For imaging GFP-labeled red blood cells, only the blue illumination (3.2 mW mm^−2^) was used.

The DMD provides flexible, programmable control over both spectral and temporal excitation. Spatial patterns can be defined to alternate between blue and lime illumination, enable single-channel constant excitation, or generate mixed dual-color patterns. By modulating the pattern sequence and duty cycle, the relative intensity of each LED channel can be dynamically tuned to achieve gradual or stepwise illumination balance. This programmability supports precise temporal encoding of dual-channel excitation and facilitates advanced illumination strategies such as alternating-frame or ramped-intensity excitation. The DMD switching frequency sets the upper bound of the achievable imaging rate and can be adapted to match the dynamics of different samples. For phantom experiments, the switching interval was 400 *µ*s—comparable to the pixel latency of the event-based camera (EVK4, Prophesee). For bead-flow assays, the interval was 500 *µ*s, yielding an effective dual-channel rate of 1 kHz. For zebrafish imaging, a 2 ms switching interval was used (250 Hz dual-channel rate), sufficient to capture the biological dynamics of interest.

After leaving the illumination module, the combined excitation beams are reflected by a dual-band dichroic mirror (59009bs, Chroma) and focused onto the sample by a 4×/0.20 NA objective (MRD70040, Nikon). The emitted fluorescence is collected by the same objective and transmitted back through the dichroic. A dual-band emission filter set (ET522/43m and ET632/65m, Chroma) spectrally separates the green and red fluorescence signals.

To engineer the system’s point-spread function (PSF), a 4*f* relay system follows the tube lens, with an SLM (PLUTO-2.1, HOLOEYE) placed at the Fourier plane for wavefront modulation. The custom phase mask—optimized using our genetic-algorithm approach [28]—extends the DOF by more than tenfold while preserving lateral resolution and is shared across both spectral channels for consistent depth response. The engineered fluorescence is then relayed to the event-based camera for high-speed acquisition and post-processing. Sensor noise was suppressed using an outlier-based event denoising algorithm (SI Appendix, Fig. S3).

### Fluorescent microsphere sample preparation

To fabricate the fluorescent phantom, 2 *µ*L of green fluorescent microspheres (2 *µ*m diameter, 1% concentration; Fluoro-Max Dyed Green Aqueous Particles) and 2 *µ*L of red fluorescent microspheres of the same type were each mixed into 2 mL of clear resin (Formlabs, RS-F2-GPCL-04) in separate tubes. The mixtures were homogenized using an ultrasonic probe sonicator (Fisherbrand Model 50 Sonic Dismembrator) to ensure uniform microsphere dispersion. Equal portions of the green and red suspensions were then combined and dispensed into a rectangular mold with a depth of 200 *µ*m. A glass slide was gently placed atop the mold to flatten the surface, and the resin was cured under UV illumination to form a solid, optically transparent phantom. After curing, the slide was removed, yielding a uniform dual-color fluorescent phantom with a precisely defined thickness of 200 *µ*m for quantitative optical characterization.

For flow experiments, a dual-color microsphere suspension was prepared by pipetting 2 *µ*L each of green and red fluorescent microspheres (2 *µ*m diameter, 1% concentration) into 2 mL of distilled water, followed by ultrasonic homogenization to minimize aggregation. A custom microfluidic channel was assembled using glass slides and coverslips. Narrow coverslip strips (100 *µ*m thick) were stacked in two layers along the long edges of a glass slide to define a channel height of approximately 200 *µ*m. A clean coverslip was placed on top and sealed along the edges to complete the flow chamber.

To aid alignment during imaging, a sparse layer of microspheres was immobilized on the top and bottom channel surfaces, providing static reference markers. During experiments, the microsphere suspension was introduced at one end of the channel, and continuous flow was maintained by pipetting fresh solution into the inlet while placing an absorbent tissue at the outlet to draw fluid by capillary action. This configuration provided a stable, gravity-driven flow.

### Design of the phase mask on the SLM

A genetic algorithm (GA) was employed to design a phase mask that extends the DOF, following the framework described in [28]. The mask’s phase profile was constructed from a basis of three phase functions—axicon, defocus, and spherical aberration—commonly used in EDOF PSF engineering. A wave-optics forward model simulated how each candidate mask modified the PSF across the axial range of interest. The merit function jointly evaluated performance in both spectral channels, maximizing PSF sharpness and peak stability across a target axial range while penalizing undesired intensity variations outside that range.

The GA refined the mask design iteratively using a “queen bee” strategy, in which the best-performing candidates were preserved between generations while crossover and mutation operations introduced new phase variations to explore the parameter space. Convergence was typically achieved within 10 generations, each containing ∼60 candidates, requiring approximately 80 minutes of computation. The resulting optimized mask provided ∼10× improvement in DOF relative to the unmodified PSF while maintaining high lateral resolution. The optimized phase pattern was implemented on an SLM.

### Dual-channel motion correction and local motion quantification

Dual-color fluorescence image stacks were split into individual red (R) and green (G) channels. Rigid-body registration was performed on the G channel using MultiStackReg (Fiji/ImageJ) to correct for global translational and rotational motion across frames. The resulting transformation matrices were applied to the R channel to ensure pixel-level spatial correspondence between channels, and the registered stacks were merged to generate a globally motion-corrected dual-color sequence. Local cellular displacements were quantified using TrackMate (Fiji/ImageJ), with detection and tracking performed independently on each channel.

### Analysis of blood flow in zebrafish

Temporal frames were reconstructed from the recorded event streams for quantitative flow analysis. Particle image velocimetry (PIV) was performed in ImageJ to calculate blood cell displacement between consecutive frames, and velocities were obtained by dividing the measured displacement by the inter-frame time interval. A multi-pass PIV algorithm was used with an interrogation window size of 32 pixels and 50% overlap to balance spatial precision and robustness. For each vessel, a short line segment along the flow direction was selected, and the mean velocity within this region was reported as the representative measurement. To generate kymographs, temporal frames recorded over a 3s interval were imported into ImageJ, and the Reslice function was applied along the line segments indicated by arrows in Fig. 6B.

### Zebrafish husbandry

Zebrafish husbandry and handling were performed in the aquatic facility at Boston University Chobanian & Avedisian School of Medicine under approved protocols from the Institutional Animal Care and Use Committee (IACUC). The Tg(d*µ*n:MYCN; d*µ*n:mCherry) and Tg(mpo:EGFP) fish in the AB wild-type background were crossed to generate the progeny for imaging. The Tg(lcr:EGFP) fish were bred to generate embryos to track red blood cells.

## Acknowledgments

The authors acknowledge funding from National Institutes of Health (R01NS126596, R01CA215059, R01NS140967), a grant from 5022 - Chan Zuckerberg Initiative DAF, an advised fund of Silicon Valley Community Foundation, and an innovation grant from the Alex Lemonade Stand Foundation. The authors acknowledge Boston University Shared Computing Cluster for proving the computational resources.

