## Supplementary Information for "Dual-Channel Event Microscopy for Ultrafast Biological Imaging"

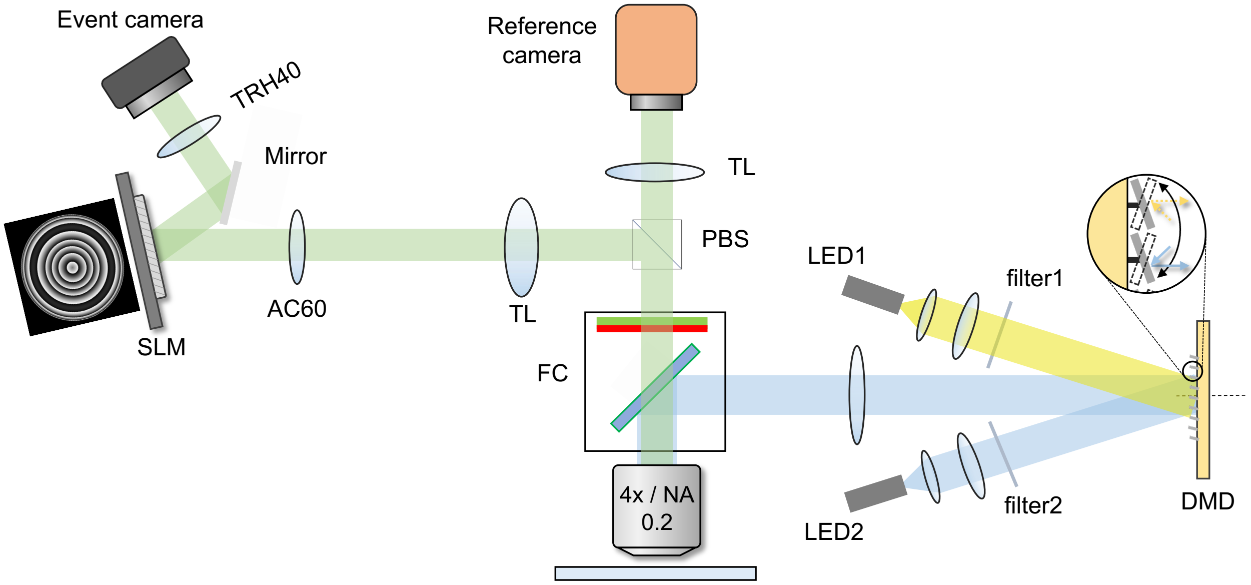


Figure S1: Schematic of the dual-channel event-based microscopy (DEM) system. A high-power blue LED (SOLIS-470C, Thorlabs) and lime LED (SOLIS-565D, Thorlabs) illuminate the digital micromirror device (DMD; DLP7000, Digital Light Innovations) at ±24° relative to the surface normal. The micromirrors toggle between ±12° at kilohertz rates to alternate blue and lime excitation, temporally multiplexing both channels within a single optical path. Each LED beam is collimated, filtered by band-pass excitation filters (ET475/35x, Thorlabs; ET575/35x, Chroma), and relayed onto the DMD to form quasi-parallel illumination. The combined excitation is reflected by a dual-band dichroic mirror (59009bs, Chroma) and focused onto the sample by a 4×/0.20 NA objective (MRD70040, Nikon). Emitted fluorescence passes back through the dichroic and dual-band emission filters (ET522/43m and ET632/65m, Chroma). A 4f relay with a spatial light modulator (SLM; PLUTO-2.1, HOLOEYE) positioned at the Fourier plane modulates the wavefront with an optimized phase mask that extends the depth of field by over tenfold while maintaining lateral resolution. The engineered fluorescence is directed to an event-based camera (EVK4, Prophesee) for high-speed acquisition and reconstruction. The DMD modulation frequency—set between 0.4 ms and 2 ms per switch depending on sample dynamics—encodes spectral information in time, enabling kilohertz-rate dual-channel imaging without mechanical switching, preserving both spatial fidelity and photon efficiency.

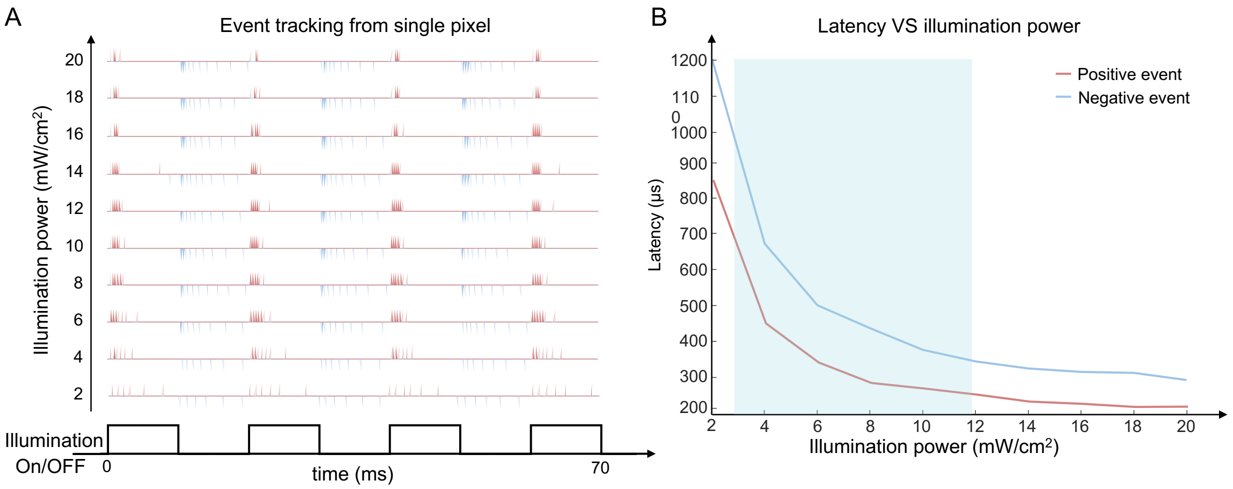


Figure S2: Characterization of event-camera pixel latency under varying illumination power. A uniform fluorescent slide (Thorlabs, FSK2) was used to evaluate pixel-level response latency at different illumination intensities. The fluorescence brightness was adjusted to approximate the emission levels of biological samples and fluorescent beads used in our experiments. The slide was fixed in the field of view, and the DMD illumination was modulated at 50Hz to generate periodic blinking excitation. (A) Event streams recorded from a single pixel show the temporal response of the sensor under different illumination powers, where each pulse corresponds to a single event. (B) Measured pixel latency as a function of illumination power for both positive (red) and negative (blue) events. Positive events, triggered by brightness increments, exhibit faster response times. In contrast, negative events, which correspond to brightness decrements, respond more slowly. As illumination power increases, both event types display markedly reduced latency, converging to ∼200µs at high intensities. The shaded blue region denotes the illumination range corresponding to fluorescence brightness levels typical of our experimental samples.

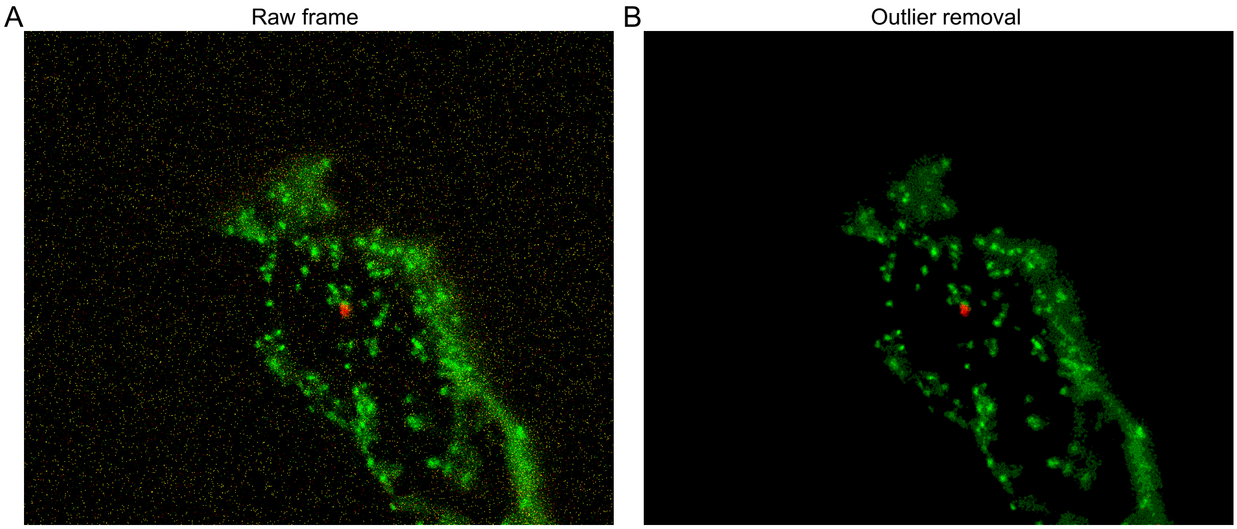


Figure S3: Event-frame denoising using outlier filtering. (A) Raw integrated event frame acquired from a freely moving zebrafish with GFP-labeled neutrophils and mCherry-labeled premalignant tumors. The raw event frame exhibits numerous isolated spurious events. (B) Denoised event frame obtained after applying an outlier removal filter in ImageJ. The filter radius was set to 2 pixels, removing isolated noisy events that deviate significantly from their local neighborhood while preserving the morphology of cellular structures and fine spatial details. This outlier-based denoising procedure effectively suppresses event-level sensor noise, improving signal-to-noise ratio and enabling clearer visualization of biologically relevant motion-encoded fluorescence signals.

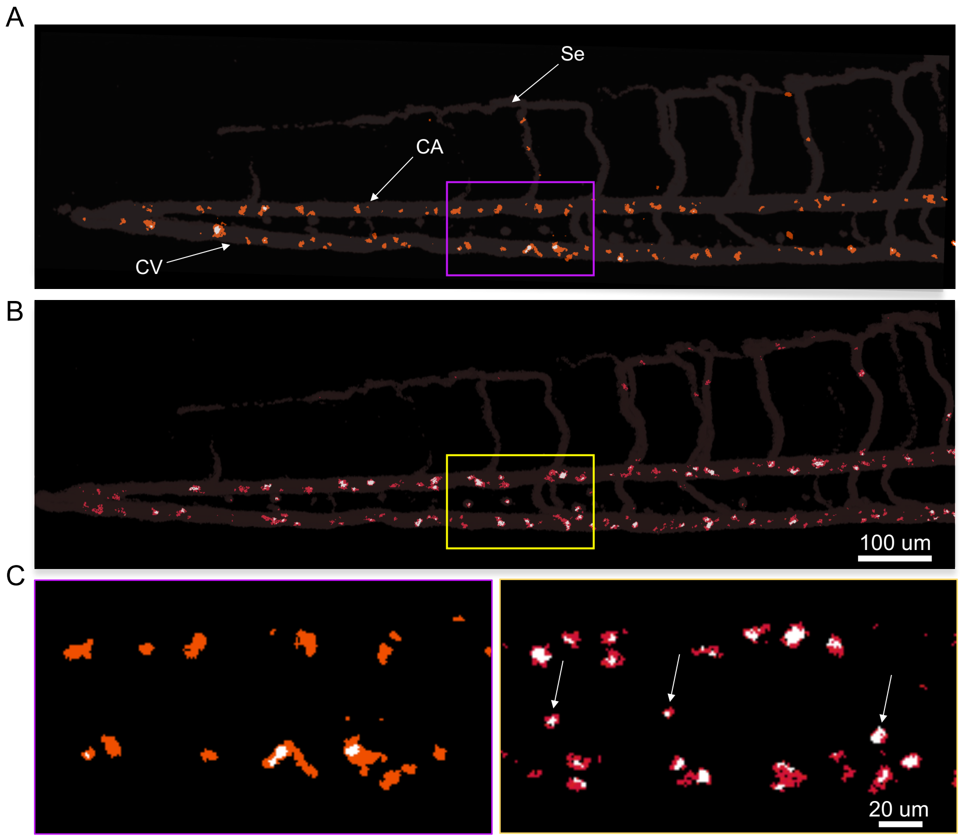


Figure S4: Comparison of blood flow imaging in zebrafish under constant and DMD-modulated illumination. (A) Reconstructed image of a zebrafish (4 dpf) with GFP-labeled red blood cells under constant blue LED illumination. Dynamic blood cells in the caudal artery (CA), caudal vein (CV), and segmental vessels (Se) are clearly visualized, while static cells remain undetected. (B) Reconstructed image of the same region acquired with temporally modulated illumination controlled by the DMD (switching interval: 2 ms). The modulation enhances the system's sensitivity to both dynamic and static fluorescence signals. (C) Zoomed-in comparison of corresponding regions (outlined in A and B). Static red blood cells, invisible under constant illumination, become clearly distinguishable when using DMD-modulated excitation, demonstrating the system's ability to capture both stationary and moving components in a single recording.

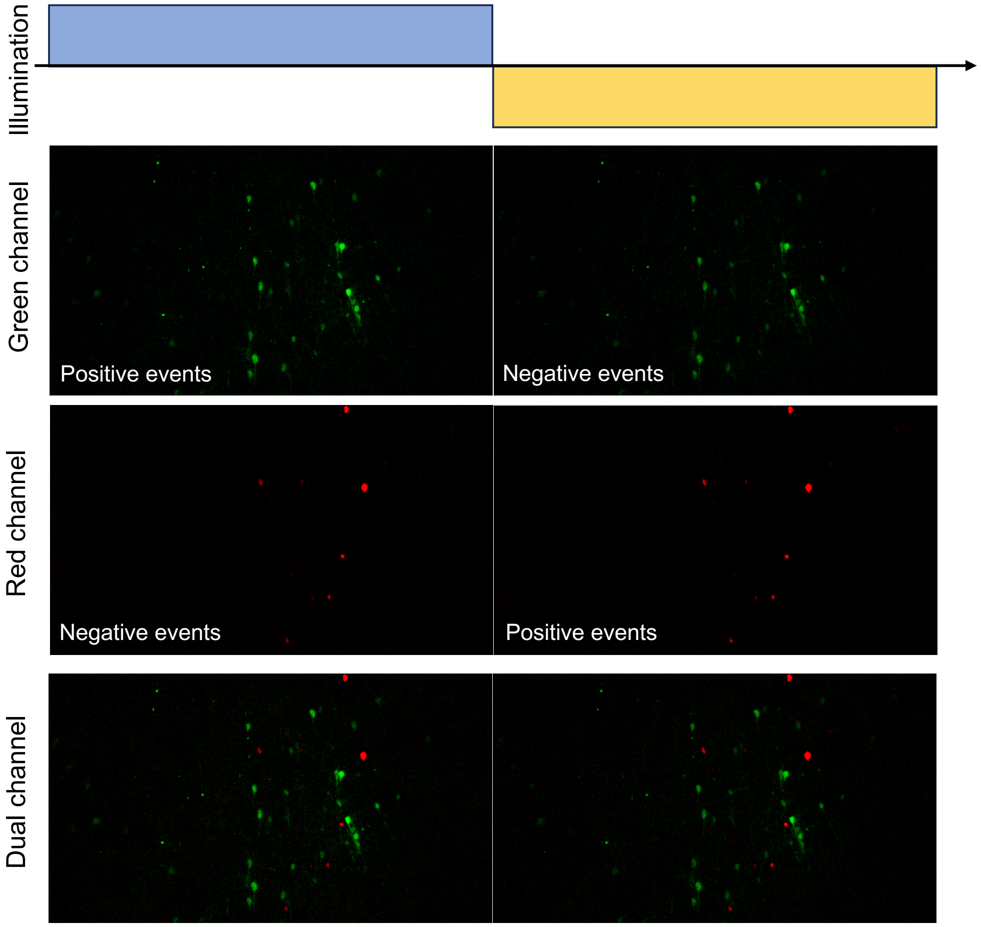


Figure S5: Dual-color imaging of a static brain slice under alternating illumination. A fixed brain slice with neurons labeled by tdTomato (red) and eGFP (green) was imaged under alternating excitation provided by the DMD. The DMD operated at 50 Hz, corresponding to a 10 ms illumination window for each color channel. The sample remained static during acquisition. Positive and negative event frames were separated for each channel to illustrate the temporal encoding mechanism. In the second and third rows, single-channel acquisitions for the green and red channels were recorded separately, where positive and negative events correspond to the LED being turned on and off during each illumination cycle. Thus, positive and negative events in each channel are recorded sequentially, serving as single-color controls. In contrast, the fourth row shows the dual-color acquisition, in which alternating blue (GFP excitation) and lime (tdTomato excitation) illumination generates temporally multiplexed positive and negative events from both channels. When illumination switches from blue to lime, neurons that turn dark produce negative events, while those newly excited generate positive events. In the green channel, positive events arise from fluorescence activated by blue illumination, whereas negative events occur as the excitation transitions to lime. Conversely, in the red channel, positive events appear under lime excitation and negative events when switching back to blue. By integrating positive and negative events across alternating illumination periods, a composite dual-color image can be reconstructed at an effective frame rate of 100 Hz, even though the illumination alternates at 50 Hz. This temporal multiplexing strategy enables high-speed, crosstalk-free dual-color imaging—static neurons can be faithfully captured under switching illumination.
